# Unlocking the microblogging potential for science and medicine

**DOI:** 10.1101/2022.04.22.488804

**Authors:** Aditya Sarkar, Augustin Giros, Louis Mockly, Jaden Moore, Andrew Moore, Anish Nagareddy, Boyang Fu, Andrada Fiscutean, Karishma Chhugani, Nicholas Darci-Maher, Yesha M. Patel, Varuni Sarwal, Yutong Chang, Srishti Ginjala, Lana X. Garmire, Riyue Bao, Sriram Sankararaman, Rayan Chikhi, Serghei Mangul

**Affiliations:** Department of Computer Science, University of California Los Angeles, 580 Portola Plaza, Los Angeles, CA 90095, USA; Ecole Centrale, Paris, France; Department of Clinical Pharmacy, School of Pharmacy, University of Southern California, 1540 Alcazar Street, Los Angeles, CA 90033, USA; Viterbi School of Engineering, University of Southern California, 1985 Zonal Avenue, Room 713. Los Angeles, CA 90089-9121, USA; Department of Pharmacology and Pharmaceutical Sciences, School of Pharmacy, University of Southern California, 1985 Zonal Avenue, Room 713. Los Angeles, CA 90089-9121, USA; Department of Computational Medicine and Bioinformatics, University of Michigan 1600 Huron Parkway, Ann Arbor 48105; UPMC Hillman Cancer Center, University of Pittsburgh Department of Medicine Pittsburgh, PA 15232; Department of Computational Biology, Institut Pasteur & CNRS, Paris, France; Indian Institute of Technology, Mandi; University of Bucharest, Bucharest, Romania

## Abstract

Microblogging platform Twitter allows researchers to showcase their work, receive constructive feedback, find jobs, and build scientific collaborations. While existing literature has analyzed the benefits of Twitter in the development and distribution of scientific knowledge, most of the studies only took into account a limited number of researchers, which affected the generalizability of derived results. Our study analyzed the activity of 6,000 biomedical scientists on Twitter using data-driven approaches, a third of whom were female. Furthermore, we estimated that up to a quarter of the members of the scientific community are engaged on Twitter. While the number of male scientists joining the microblogging platform every year has decreased, the number of female scientists has remained roughly the same. Scientists are very selective in who they are following as compared to the general public. We also found that the type of tweets and retweets one posts may affect the number of followers, specifically, that a moderate to high level of professionalism and a high level of positivity is correlated with followers count. Moreover, female scientists send fewer negative tweets as compared to male scientists (31.1% for females and 34.7% for males). Our analysis could provide insights and launch a conversation on the advantages and limitations of using Twitter for disseminating scientific information and engaging in constructive discussion and collaborations within the scientific community.

## Introduction

Twitter is one of the most popular microblogging and social networking platforms allowing users to post, retweet, comment, and engage in discussions^1^. In combination with preprint services widely used by the biomedical community, it promises to mitigate barriers in terms of accessing information, leading to a major impact especially during the early stages of one’s scientific career^2^, when it is hard to find opportunities. Recent studies^3^ show that dissemination of research work over social media can be used as an academic metric, and one way to quantify it is by using the tweets. Twitter provides an effective platform for scientists to interact and establish novel collaborations with peers, yet its full potential might not be accessed. For instance, when tangible networking opportunities arise, including during virtual conferences, the number of Tweets posted by scientists might even go down, instead of up.

Existing analyses^5–10^ of the activity of scientists on Twitter have shown a relationship between Twitter mentions and article citations, as well as benefits of Twitter in the development and distribution of scientific knowledge, and the content that researchers tweet. However, most existing studies are limited by sample size and have only analyzed a limited number of researchers, potentially compromising generalizability of reported results. For instance, a recent survey^1^ tried to understand the ways scientists interact on Twitter and concluded that the platform keeps researchers updated on new preprints, publications, innovations, opportunities etc. It also helps them to self-promote their work, makes them feel connected to the scientific community and helps them to interact with like-minded scientists. The survey was only able to reach a limited number of researchers (less than 100) who agreed to participate, thus being unable to fully capture the more general trends of using the social platform by a biomedical community. In addition, it is not known how factors such as the length of time scientists are using Twitter, their current career status, and their following practices can affect the followers count. Furthermore, how scientists are following or getting followed, and their activity on the platform are also unclear.

Our analysis focuses on these trends, and overcomes a key limitation found in previous studies by analyzing substantially larger data accrued for around six thousand scientists. Our ultimate goal is to quantify the important characteristics of scientific Twitter including components such as the fraction of scientists using Twitter, the length of time they spend using the platform, their current career status, and how they are followed. In addition to that, we compare the time scientists spend on Twitter in regards to the career stage.

## Data Collection

To analyze the Twitter activity of biomedical researchers, we needed to collect their Twitter accounts. To get started, we first filtered out the information of 167,000 scientists who have published at least five publications. Then, web scraping techniques were used to identify the corresponding Twitter accounts of each of the 167,000 researchers based on first and last names. As a result, we obtained 873,000 candidate Twitter handles matching the names of scientists. Next, the authors’ Twitter data, i.e., number of followers, number of people they are following, status count, description, their tweets etc., were collected using the Tweepy API. Using the extracted Twitter descriptions as a filter, we conservatively extracted all the Twitter IDs belonging to the scientific field by finding keywords related to academic titles in their bio, for example, “professor,” “postdoc,” “Ph.D.”, etc. Duplicate accounts were resolved by picking up the first occurrence of the Twitter account over a Google search of the scientist’s name. At the end of the process, we were left with a total of 6,124 scientists. Next, we removed those scientists who hadn’t tweeted a single tweet, which consisted of 5.32% of 6,142. Thus, the number was reduced to 5,798 scientists, on whom we performed all of our analyses.

To validate the proposed approach of matching names and Twitter IDs, we prepared a gold-standard data by randomly selecting 200 researchers and then manually matching their Twitter profiles with the data collected, to check if the accounts did indeed belong to the scientist. As a result, we got an accuracy of 0.98, with a specificity of 0.994, a sensitivity of 0.57 and a precision of 0.8. Additionally, we used Naive Bayes Classifier to predict the gender of 5798 scientists using the first names. Out of these scientists, 3,787 (65.3%) were predicted to be male and 2,011 (34.68%) to be female.

### Up to a quarter of the members of the scientific community are engaged on Twitter

We first assessed the fraction of biomedical researchers who utilize Twitter by estimating the minimum and maximum number of scientists using the platform. A minimum value is given by the number of accounts that are correctly predicted as scientists, estimated through the accuracy of our filters. Since our filters were able to correctly predict 98 out of 100 scientists, we consider that 98% of the 5,798 twitter accounts collected, which are associated with 5,682 people, are actually scientists. This represents 3.402% of the initial pool of 167,000 scientists. Next, to estimate the maximum number of scientists, we disable the filters in our methods otherwise allowing us to effectively remove accounts of non-scientists based on the keyword of profile description. Our filters conservatively remove accounts with non-scientific keywords in description. If we generously consider that all of these accounts belong to scientists, we estimate the maximum number of scientists using Twitter to be 37,302 or 22.3% of the total 167,000, in line with the share of U.S. adults using Twitter, estimated to be at around 23%^11^. Therefore, we estimate that 3.4-22.3% of biomedical scientists are active on Twitter.

We further observed the trend of scientists joining Twitter over the years and we found that overall, the number of scientists joining the platform has decreased in recent years. We analyzed the number of scientists who joined 10 years back, 4 years back and 2 years back, and we noticed that the numbers have gone down. 2 years back, only 528 scientists from our panel joined Twitter while 10 years back, 960 scientists joined the platform. However, while the number of male researchers joining the platform has been going down over the years, the number of female researchers has remained relatively constant. Two years back, around 300 males joined the platform, while ten years back, around 700 males joined the platform. By comparison, 210 females joined 2 years back and 285 females joined 10 years back.

### Effect of academic status on number of followers

In order to explain how we calculated dependency between variables, we will briefly explain the concept of information content, also known as Shannon entropy. Information content or Shannon entropy of a random variable is defined as the level of uncertainty present in the random variable’s possible outcomes. Further mutual information can be used to explain how much information content about the dependent variable can be explained by independent variables. We calculated mutual information between independent variables, which we have taken as gender, academic status, the number of following, count of tweets and retweets, and the dependent variable, which we took as follower count. This is the same as finding mutual information between the dependent variables and independent variables. Similar analysis was performed for different stratifications of data, where the stratification variables were gender, race and academic profession. Considering the follower count as the dependent variable and having the following count, academic profession, number of days on Twitter, status count (total number of posts), and gender as independent variables, we find that overall, the main explanation of the number of followers was being provided by number of days on Twitter, status count, number of followers and number of following. There is very little dependence of gender and academic profession on follower count.

We further analyzed the trends of followers and following count on Twitter. A typical scientist from our panel has 211 following and 176 followers, which means that the scientist follows 211 users and is followed back by 176 users. Twitter has no restrictions on following accounts, however, researchers tend to be selective who they follow. On average, they tend to follow fewer users than they are followed by, with 70% of scientists having the followers by the following ratio of 1.5. We also found that 26% of scientists had very few followers, but were following a lot of other users which made followers by the following ratio to less than 0.5. On the other hand, 23% scientists had many followers but were not following other people back to make followers by the following ratio more than 2.

We observed a significant difference in the number of followers across the professional ranking (Anova test, F-statistic=3.097, p-value=0.025). Professors have the highest median value of number of followers (309), while post-doctorates have the lowest (121), even lower than graduate students (143). We also noticed that professors and scientists who fall into the “other researchers” category follow much fewer scientists than they are followed by (followers by following ratio = 1.014). Around 53.3% of the professors have more followers than following. Out of this, 13.5% are females and 86.5% are males. However, this trend is not shown by doctorates and post-doctorates, as around 75.4% of doctorates and post-doctorates have more following than followers.

### Activity of scientists on Twitter

We also analyzed the content of tweets and retweets posted by scientists on the platform in a subjective manner. We found, through sentiment analysis (where we analyzed the subjective information of the tweets), that on an average, scientists send negative tweets 33.5% of the time, with positive tweets accounting for 64%, the remaining being neutral tweets (2.5%). Female scientists send fewer negative tweets as compared to male scientists (31.1% for females and 34.7% for males). On an average, scientists send personal tweets 75.6% of the time and professional tweets 17% of the time. Among academic professions, postdoctorates send positive tweets for 70% of the time which is the maximum, while scientists in other professions send it for 63-64% of the time.

We then analyzed the number of followers scientists have across levels of positivity of tweets. The positivity of tweets was measured between 0 and 1, with the range being divided into 5 equal sections. The conclusion was that the more positive tweets a scientist posted on Twitter, the more followers they had. Similarly we analyzed the number of followers of scientists across levels of professionalism in their tweets. The range of professionalism was again measured between 0 to 1, divided into 5 sections. We found that if a scientist posted a high number of professional tweets, the number of followers tended to decrease. On the other hand, if the proportion of professionalism is between 0 to 0.6, then the median number of followers are almost the same. From the range of 0.6 to 1, the median number of followers is falling. We find that the professionalism proportion of 0.4 to 0.6 and positivity proportion of 0.6 to 0.8 is ideal for increasing followers count.

## Discussion

Twitter provides scientists the opportunity to showcase their work and engage in conversations with researchers across the world. It can be powerful in mitigating barriers in access to information, leading to a major impact especially during the early stages of one’s scientific career^2^, when finding career opportunities can be a struggle. In this context, our analysis aims to help researchers to unlock the potential of Twitter. The study has quantified how factors such as the length of time scientists are using Twitter, their current career status, gender, and following count affect their followers count.

The trends of how the scientists are following or getting followed, and their activity on the platform were also analyzed. We observed a high dependence of following count and activity of scientists on followers count, which indicates that researchers have to be active on the platform in order to increase their followers count. In addition to that, we noticed that professors tend to follow fewer users but are followed more likely because to advance their career, PhD students and postdoctorates tend to follow more people than they are followed. This enables them to improve their connections and thereby their chances of accessing career opportunities.

There are certain limitations to our study. While predicting gender, we relied on machine learning techniques. Despite using these measures for our predictions, we recognise the accuracy of machine learning methods used to infer gender has yet to be validated and thus should be used with caution. Another limitation associated with the Twitter platform is that improper usage of Twitter can be detrimental to science and even have a negative impact on mental health. The number of scientists using Twitter seems to be lower compared to the general population.

Our analysis gives arguments that scientific Twitter provides an effective platform for researchers to disseminate ideas, establish new collaborations and discuss scientific ideas^9,12^. The ability to leverage Twitter by the scientific community is an essential skill the effects of which will not only help accelerate the spread of broad scientific ideas among diverse communities, but also gain communal feedback. This will provide an effective means to substantially improve scientific work, which can be used in addition to or instead of traditional peer review methods^4,13,14^.

Engaging in field-specific discussions on Twitter allows researchers to develop and expand their professional network, launch new research projects and collaborations, and receive help from the community at various stages of their projects. It aids researchers in presenting their work and supports international scientists. For example, the Twitter account (@Sci_for_Ukraine) was created to support scholars and students affected by Russia’s war in Ukraine; it gained 7,900 followers in 40 days. Together with the website and other social media accounts, it is curated by an international community of volunteering scholars.

Our analysis indicated that gender backgrounds had some impact on the Twitter parameters. Future work is needed to fully understand and interpret the differences we have noted across gender and ancestry groups. We also hope the results presented here will encourage researchers to join Twitter and become fully engaged on social media for sharing their ideas and contribution across the global scientific community. We believe social media can be effectively used as a harbor of scientific knowledge to widely disseminate novel ideas and innovations among the global research community beyond geographical and administrative boundaries.

## ACKNOWLEDGEMENTS

We thank Jeremy Rotman for his assistance in data analysis. Our paper is dedicated to all freedom-loving people around the world. We would like to thank Sanita Reinsone (@sanitare) for helpful discussions.

## DECLARATIONS

### Data and Material Availability

All supporting material for this analysis can be found at: https://github.com/Mangul-Lab-USC/Twitter-editorial

### Contributions

SM and RC conceived of the presented idea. NDM prepared the datasets. AS conducted the major analysis of the datasets and wrote the scripts for initial data preprocessing and analysis. YC assisted in curating the data. LM and AG contributed to data analysis. RC, AS, KC, VS, AF and SM contributed to the writing of the manuscript. All authors discussed the text and commented on the manuscript. All authors read and approved the final manuscript.

